## SupplementaryMaterial for "Alcohol consumption and mate choice in UK Biobank: comparing observational and Mendelian randomization estimates"

**Supplementary Methods**

Assuming a population of unrelated individuals (1000 males *M* and 1000 females *F*) where a phenotype *P* is assorted on (individuals more phenotypically similar for *P* are more likely to pair-up).

If *P* $\sim N(0,1)$ and is influenced by genetic factors $G\sim N(0,1)$ such that:

$$Cov\left( G,P \right)=\left( \begin{matrix} 1 & \sqrt{h^{2}} \\ \sqrt{h^{2}} & 1 \end{matrix} \right)w\mathrm{here}h^{2} is the proportion of variation in P explained by G$$

$It then follows that with assortment on P such that in each male female pair Cor\left( M_{P},F_{P} \right)=C$, that the expected Mendelian randomization and genetic correlation estimates capturing spousal assortment are equivalent to:

$$Cor\left( M_{P},F_{G} \right)=Cor\left( M_{G},F_{P} \right)=\frac{Cov(M_{G}, F_{P})}{\sigma_{M_{G}}\sigma_{F_{P}}}=Cov\left( M_{G}, F_{P} \right)=\left( \begin{matrix} 1 & \sqrt{C}\sqrt{h^{2}} \\ \sqrt{C}\sqrt{1-h^{2}} & 1 \end{matrix} \right)$$

$$=C*\sqrt{h^{2}}*\sqrt{{1-h}^{2}}$$

$$Cor\left( M_{G},F_{G} \right)=\frac{Cov\left( M_{G}, F_{G} \right)}{\sigma_{M_{G}}\sigma_{F_{G}}}=Cov\left( M_{G}, F_{G} \right)=\left( \begin{matrix} 1 & \sqrt{C}\sqrt{h^{2}} \\ \sqrt{C}\sqrt{h^{2}} & 1 \end{matrix} \right)$$

=$C*h^{2}$

Furthermore, we used simulated data to confirm that same-trait spousal Mendelian randomization (GxP) and genetic correlation (GxG) estimates are influenced by the heritability (h^2^) of the trait and the degree of phenotypic assortment C. We used a range of parameters: (h^2^: 0.05, 0.1, 0.15, 0.2, 0.3, 0.4, 0.5) & (C: 0.1, 0.2, 0.3, 0.4, 0.5). 1000 simulations were run for each model with the mean correlation estimate reported and these results were consistent with the derived formulae.

Simulation results:

| **Heritability of P** | **Degree of assortment: Phenotypic correlation C** | **Correlation (GxG, GxP):**  **Mean of 1000 simulations** |
| --- | --- | --- |
| 0.05 | 0.1 | 0.005, 0.023 |
|  | 0.2 | 0.008, 0.046 |
|  | 0.3 | 0.015, 0.067 |
|  | 0.4 | 0.021, 0.090 |
|  | 0.5 | 0.026, 0.110 |
| 0.1 | 0.1 | 0.010, 0.033 |
|  | 0.2 | 0.020, 0.062 |
|  | 0.3 | 0.030, 0.092 |
|  | 0.4 | 0.039, 0.126 |
|  | 0.5 | 0.050, 0.157 |
| 0.15 | 0.1 | 0.015, 0.038 |
|  | 0.2 | 0.029, 0.078 |
|  | 0.3 | 0.045, 0.116 |
|  | 0.4 | 0.059, 0.156 |
|  | 0.5 | 0.074, 0.194 |
| 0.2 | 0.1 | 0.021, 0.046 |
|  | 0.2 | 0.040, 0.089 |
|  | 0.3 | 0.060, 0.133 |
|  | 0.4 | 0.079, 0.177 |
|  | 0.5 | 0.101, 0.225 |
| 0.3 | 0.1 | 0.030, 0.054 |
|  | 0.2 | 0.059, 0.108 |
|  | 0.3 | 0.088, 0.164 |
|  | 0.4 | 0.121, 0.220 |
|  | 0.5 | 0.150, 0.275 |
| 0.4 | 0.1 | 0.040, 0.064 |
|  | 0.2 | 0.078, 0.127 |
|  | 0.3 | 0.119, 0.190 |
|  | 0.4 | 0.158, 0.251 |
|  | 0.5 | 0.198, 0.315 |
| 0.5 | 0.1 | 0.049, 0.070 |
|  | 0.2 | 0.099, 0.141 |
|  | 0.3 | 0.149, 0.212 |
|  | 0.4 | 0.201, 0.282 |
|  | 0.5 | 0.250, 0.353 |

**Supplementary Table 1: Information on alcohol questionnaire variables**

| **Alcohol variable** | **Median (Q1, Q3); Max** |
| --- | --- |
| *Alcoholic units a week (N=95,059)^1^:*  Spirit measures  Glasses of white wine  Glasses of red wine  Glasses of fortified wine  Pints of beer or cider | 12 (1, 24); 312  0 (0,1); 200  0 (0,2); 80  1 (0,4); 90 0 (0,0); 84  0 (0,2); 72 |
| *Current drinking status (N= 95,059):*  Never: N (%)  Previous: N (%)  Current: N (%) | 2669 (2.8%)  2630 (2.8%)  89760 (94.4%) |
| *Current alcohol intake frequency (N= 95,059):*  Never: N (%)  Special occasions only: N (%)  One to three times a month: N (%)  Once or twice a week: N (%)  Three or four times a week: N (%)  Daily or almost daily: N (%) | 5299 (5.6%)  8583 (9.0%)  9497 (10.0%)  25253 (26.6%)  25222 (26.5%)  21205 (22.3%) |

^1 Self-report non or former drinkers had values imputed to 0 for relevant variables.^

**Supplementary Table 2:** Weekly alcohol consumption per rs1229984 genotype

|  | **385,287 individuals of European descent** | | | **337,114 individuals of White British descent** | | |
| --- | --- | --- | --- | --- | --- | --- |
|  | **CC (N=363,036)**  **Median (Q1, Q3)** | **TC (N=20,146)**  **Median (Q1, Q3)** | **TT (N=569)**  **Median (Q1, Q3)** | **CC (N=320,699)**  **Median (Q1, Q3)** | **TC (N=14,728)**  **Median (Q1, Q3)** | **TT (N=188)**  **Median (Q1, Q3)** |
| **Weekly alcohol consumption (units)** | 12.0 (0, 24) | 7.0 (0, 17.5) | 2.0 (0, 13) | 12.0 (0, 24.0) | 8.0 (0, 18) | 4.3 (0, 18.1) |

**Supplementary Table 3:** Association of rs1229984 with genetic principal components and birth coordinates

|  | **CC (%)** | **TC (%)** | **TT (%)** | **Hardy-Weinberg:** Chi^2^, P-value |
| --- | --- | --- | --- | --- |
| **385,287 individuals of European descent** | 364,520 (94.6%) | 20,194 (5.2%) | 573 (0.1%) | 274.7, <10^-16^ |
| **337,114 individuals of White British descent** | 322,183 (95.6%) | 14,743 (4.4%) | 188 (<0.1%) | 2.0, 0.16 |

**Supplementary Table 4:** Association of rs1229984 with genetic principal components and birth coordinates

| **Principal components** | **385,287 individuals of European descent** | | **337,114 individuals of White British descent** | |
| --- | --- | --- | --- | --- |
|  | **P-value** | | **P-value** | |
| PC1 | <10^-16^ | | <10^-16^ | |
| PC2 | <10^-16^ | | <10^-16^ | |
| PC3 | <10^-16^ | | 0.90 | |
| PC4 | <10^-16^ | | <10^-16^ | |
| PC5 | 0.07 | | <10^-16^ | |
| PC6 | 0.04 | | 0.00022 | |
| PC7 | <10^-16^ | | 0.074 | |
| PC8 | 0.07 | | 0.11 | |
| PC9 | 0.86 | | 0.11 | |
| PC10 | 1.53x10^-9^ | | 0.00070 | |
| **Birth-coordinates (units)** | **Beta (95% C.I.)^1^** | **P-value** | **Beta (95% C.I.)^1^** | **P-value** |
| North-South Axis (miles north) | 24.6 (22.2, 27.0) | <10^-16^ | 19.4 (16.7, 22.0) | <10^-16^ |
| East-West Axis (miles east) | -13.3 (-14.5, -12.1) | <10^-16^ | -10.3 (-11.7, -9.0) | <10^-16^ |

^1 Per additional major allele (associated with increased alcohol consumption)^

**Supplementary Table 5:** Association of self-reported weekly alcohol consumption with genetic principal components and birth coordinates

| **Principal components** | **385,287 individuals of European descent** | | **337,114 individuals of White British descent** | |
| --- | --- | --- | --- | --- |
|  | **P-value** | | **P-value** | |
| PC1 | <10^-16^ | | 3.3x10^-11^ | |
| PC2 | <10^-16^ | | 0.57 | |
| PC3 | <10^-16^ | | 3.2x10^-6^ | |
| PC4 | <10^-16^ | | <10^-16^ | |
| PC5 | <10^-16^ | | <10^-16^ | |
| PC6 | <10^-16^ | | 0.36 | |
| PC7 | 0.18 | | 1.1x10^-5^ | |
| PC8 | 2.2x10^-10^ | | 0.0094 | |
| PC9 | <10^-16^ | | 2.9x10^-10^ | |
| PC10 | <10^-16^ | | 1.8x10^-12^ | |
| **Birth-coordinates (units)** | **Beta (95% C.I.)^1^** | **P-value** | **Beta (95% C.I.)^1^** | **P-value** |
| North-South Axis (kilometres north) | 0.19 (0.16, 0.22) | <10^-16^ | 0.18 (0.15, 0.21) | <10^-16^ |
| East-West Axis (kilometres east) | -0.03 (-0.04, -0.02) | 7.1x10^-5^ | -0.03 (-0.04, -0.01) | 4.4x10^-4^ |

^1 Per 1 unit increase in weekly alcohol consumption^

**Supplementary Table 6:** Spousal birth proximity and association with birth coordinates and principal components

|  | **Complete spouse sample**  **(N~47,549)** | | | | **Spouses born within 100km (N~28,580)** | | | | **Spouses born more than 100km apart (N~13,746)** | | | |
| --- | --- | --- | --- | --- | --- | --- | --- | --- | --- | --- | --- | --- |
|  | **Spouse alcohol consumption difference per week (units)** | | **Spousal rs1229984 genotype differences** | | **Spouse alcohol consumption difference per week (units)** | | **Spousal rs1229984 genotype differences** | | **Spouse alcohol consumption difference per week (units)** | | **Spousal rs1229984 genotype differences** | |
| **Spousal principal components difference** | **P-value** | | **P-value** | | **P-value** | | **P-value** | | **P-value** | | **P-value** | |
| PC1 | 1.22x10^-8^ | | <10^-16^ | | 0.007 | | <10^-16^ | | 0.003 | | <10^-16^ | |
| PC2 | 2.28x10^-7^ | | <10^-16^ | | 0.49 | | <10^-16^ | | 1.3x10^-4^ | | <10^-16^ | |
| PC3 | <10^-16^ | | <10^-16^ | | 3.09x10^-5^ | | <10^-16^ | | 7.63x10^-13^ | | <10^-16^ | |
| PC4 | <10^-16^ | | <10^-16^ | | 4.72x10^-16^ | | <10^-16^ | | 8.29x10^-16^ | | <10^-16^ | |
| PC5 | 9.69x10^-13^ | | 0.70 | | 4.89x10^-12^ | | 0.14 | | 5.93x10^-4^ | | 0.57 | |
| PC6 | 0.27 | | 6.88x10^-5^ | | 0.12 | | 0.010 | | 0.14 | | 2.37x10^-4^ | |
| PC7 | 0.60 | | <10^-16^ | | 0.97 | | <10^-16^ | | 0.37 | | <10^-16^ | |
| PC8 | 0.12 | | 0.15 | | 0.031 | | 0.10 | | 0.69 | | 3.57x10^-7^ | |
| PC9 | 0.15 | | 0.90 | | 0.13 | | 0.71 | | 0.16 | | 0.98 | |
| PC10 | 0.037 | | 0.10 | | 0.55 | | 0.082 | | 0.0027 | | 0.23 | |
| **Spousal Birth-coordinates difference (units)** | **Beta (95% C.I.)^1^** | **P-value** | **Beta (95% C.I.)^2^** | **P-value** | **Beta (95% C.I.)^1^** | **P-value** | **Beta (95% C.I.)^2^** | **P-value** | **Beta (95% C.I.)^1^** | **P-value** | **Beta (95% C.I.)^2^** | **P-value** |
| North-South Axis (kilometres north) | 0.18 (0.10, 0.25) | 1.3x10^-6^ | 9.3 (5.2, 13.4) | 9.4x10^-6^ | -0.003 (-0.017, 0.010) | 0.62 | 0.46 (-0.33, 1.26) | 0.25 | 0.56 (0.33, 0.78) | 1.2x10^-6^ | 25.7 (13.6, 37.8) | 3.0x10^-5^ |
| East-West Axis (kilometres east) | -0.05 (-0.09, -0.01) | 0.016 | -4.1 (1.7, 6.4) | 7.5x10^-4^ | -0.000 (-0.014, 0.013) | 0.95 | -0.75 (-1.58, 0.08) | 0.078 | -0.15 (-0.28, -0.03) | 0.019 | -10.2 (-3.4, -17.1) | 0.0034 |

^1 Per 1 unit increase in weekly alcohol consumption^

^2 Per additional major allele (associated with increased alcohol consumption)^

**Supplementary Figure 1:** Subsets of UK Biobank utilised in analyses

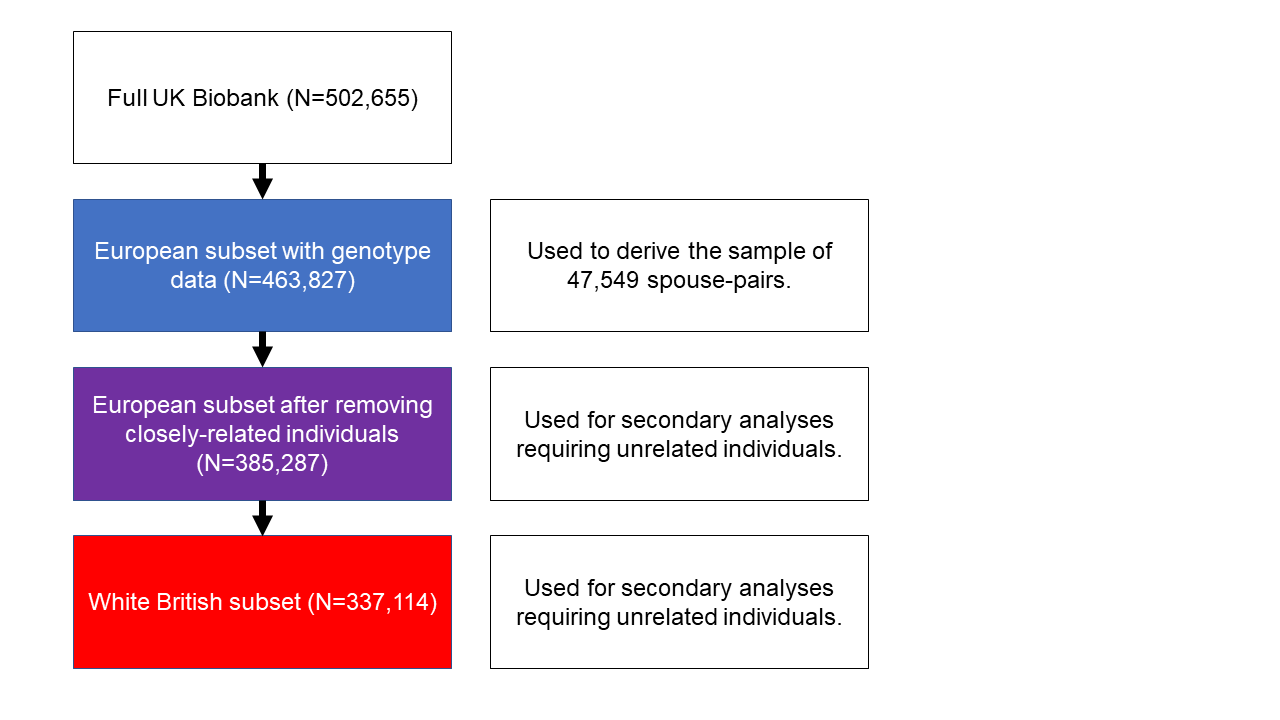
